# Microbial short-chain fatty acids regulate drug seeking and transcriptional control in a model of cocaine seeking

**DOI:** 10.1101/2023.03.22.533834

**Authors:** Katherine R. Meckel, Sierra S. Simpson, Arthur Godino, Emily G. Peck, Olivier George, Erin S. Calipari, Rebecca S. Hofford, Drew D. Kiraly

## Abstract

Cocaine use disorder represents a public health crisis with no FDA-approved medications for its treatment. A growing body of research has detailed the important connections between the brain and the resident population of bacteria in the gut, the gut microbiome in psychiatric disease models. Acute depletion of gut bacteria results in enhanced reward in a mouse cocaine place preference model, and repletion of bacterially-derived short-chain fatty acid (SCFA) metabolites reverses this effect. However, the role of the gut microbiome and its metabolites in modulating cocaine-seeking behavior after prolonged abstinence is unknown. Given that relapse prevention is the most clinically challenging issue in treating substance use disorders, studies examining the effects of microbiome manipulations in relapse-relevant models are critical. Here, Sprague-Dawley rats received either untreated water or antibiotics to deplete the gut microbiome and its metabolites. Rats were trained to self-administer cocaine and subjected to either within-session threshold testing to evaluate motivation for cocaine or 21 days of abstinence followed by a cue-induced cocaine-seeking task to model relapse behavior. Microbiome depletion did not affect cocaine acquisition on an FR1 schedule. However, microbiome-depleted subjects exhibited significantly enhanced motivation for low dose cocaine on a within-session threshold task. Similarly, microbiome depletion increased cue-induced cocaine-seeking following prolonged abstinence. In the absence of a normal microbiome, repletion of bacterially-derived SCFA metabolites reversed the behavioral and transcriptional changes associated with microbiome depletion. These findings suggest that gut bacteria, via their metabolites, are key regulators of drug-seeking behaviors, positioning the microbiome as a potential translational research target.

## Introduction

Psychostimulant use disorder is a recalcitrant neuropsychiatric condition with a relapsing-remitting course leading to profound morbidity and mortality^1^. Although understanding of the cellular and circuit-level adaptations in models of psychostimulant use disorders has advanced tremendously, translating these findings to the clinical arena has proven difficult^2,3^. As such, there are currently no FDA-approved medications for psychostimulant use disorders. This urgent need has led to increased interest in examining peripheral factors beyond the blood-brain barrier, which may serve as useful therapeutic targets or biomarkers^4,5^.

It has become increasingly clear that the resident population of bacteria within the gastrointestinal tract, collectively termed the gut microbiome, modulates brain function^6,7^. Robust literature indicates that connections between the brain and the gut microbiome modulate neuropsychiatric disease pathogenesis in animal models and clinical studies of autism, depression, anxiety, and substance use disorders^5,8–15^. Indeed, previously published work demonstrates gut microbiome depletion results in significantly altered behavioral and transcriptional responses to cocaine and opioids^11,16–21^.

Although the exact mechanisms underlying gut-brain signaling are unclear, evidence suggests that neuroactive metabolites produced by the microbiome play a signaling role in neuropsychiatric diseases^10,12,22^. The microbiome is a highly metabolically active ecosystem that produces hundreds of metabolites that are absorbed into the circulation^23–25^. Short-chain fatty acids **(SCFA)**, metabolites produced by the bacterial fermentation of dietary fiber, are key gut-brain signaling mediators^26,27^. The three primary SCFA (butyrate, acetate, and propionate) have widespread effects on brain function, rescuing effects of microbiome depletion at the behavioral and transcriptional level across neuropsychiatric disease models^27–31^. Repletion of SCFA reverses the behavioral effects of microbiome depletion in mouse models of cocaine and morphine conditioned place preference^11,16^.

Microbiome manipulations also affect animal models of opioid tolerance and hyperalgesia^20,32,33^, and more complex effects of the microbiome on cocaine place preference and sensitization^18,19^. Importantly, these early studies have focused on shorter-term experimenter-administered drug treatments. Although these studies have advanced understanding of the effects of microbiome manipulations on animal models of substance use disorders, there is a need to examine microbiome effects on models of drug self-administration, which are the most translationally-relevant animal models of substance use disorders^34^. These models provide critical information on both drug-taking and drug-seeking after abstinence, a model for drug relapse^35,36^. Relapse-type behavior arises in part due to incubation of drug craving, a phenomenon seen in both human subjects and animal models in which the desire to administer drugs increases with longer periods of abstinence^34,37,38^. Given that relapse prevention is the most difficult challenge in treating patients with psychostimulant use disorder, understanding microbiome effects in these models is critical for future translational insight.

Here, we leveraged our established antibiotic microbiome depletion model to examine the effects of microbiome manipulations on drug-seeking behavior in a translationally-relevant self-administration model for cocaine intake and relapse-like behavior. Antibiotic-induced depletion of the gut microbiome in male rats markedly alters the rewarding effects of cocaine and motivation to seek cocaine after withdrawal. Furthermore, animals with altered microbiomes have altered expression of synaptic plasticity genes following cocaine self-administration. To assess the mechanistic contribution of microbial metabolites, repletion experiments utilizing SCFA were also performed. Crucially, repletion of bacterially-derived SCFA metabolites reversed the behavioral and molecular changes associated with microbiome depletion. This study provides unprecedented mechanistic insight into the role of the microbiome and its metabolites on striatal gene expression in a translationally-relevant model of cocaine use disorder and relapse.

## Materials & Methods

### Animals

Adult male Sprague-Dawley rats (ENVIGO-Harlan, 300-320g) were pair-housed under specific-pathogen free conditions in a room with constant 22-25°C and humidity of 55%. Subjects were maintained on a reverse 12-hr light-dark cycle (lights on at 1900 hours) with food and water available *ad libitum.* All animal protocols were approved and conducted in accordance with the policies of the Institutional Animal Care and Use Committee at Mount Sinai.

### Antibiotic and Short-Chain Fatty Acid Treatments

Animals were randomly assigned and given *ad libitum* access to non-absorable antibiotics administered in the drinking water at concentrations of neomycin 2 mg/ml, vancomycin 0.5 mg/ml, bacitracin 0.5 mg/ml, and pimaricin 1.2 μg/ml (all from Fisher Scientific), in an adaption of prior studies^11,39^. For short-chain fatty acids (SCFA) experiments, the three principle bacterially-derived SCFA were dissolved in the drinking water at physiological levels (67.5mM acetate, 40mM butyrate, 25.9mM propionate, all from Sigma Aldrich) as described previously^11,40,41^. Animals were treated with antibiotics and/or SCFA for two weeks prior to any testing and were maintained on this treatment for the duration of the study.

### Cocaine Self-Administration: Acquisition

Cocaine self-administration was performed in standard operant conditioning chambers (MEDAssociates) as described previously^42^. Subjects were food restricted throughout the duration of training with 18g food/subject delivered once daily following session completion. Sessions were daily for 3 hours. Subjects were trained to respond for cocaine on the “active” lever on an FR1 schedule of reinforcement where one lever press resulted in the delivery of a 5.9s infusion of 0.8mg/kg/infusion (0.1ml) cocaine paired with concurrent illumination of the cue light located above the active lever.

### Experiment 1: Within Session Threshold Testing

Following acquisition of FR1 administration, a within session threshold test was performed to assess differences in cocaine motivation and consumption between groups. In this behavioral economics task, rats are given continuous access to cocaine while increasing the effort requirement to obtain the same amount of drug^42,43^. Importantly, this measure allows for assessment of both motivation and dose-response within a single session. Full details are included in **Supplemental Materials and Methods**.

### Experiments 2 and 3: Cue-Seeking After Abstinence

These experiments were performed in separate groups of rats from the threshold tasks. Following acquisition, rats were returned to their home cages for 21 days of abstinence. Food restriction was lifted during this abstinence period and resumed 1 day before behavioral testing. For the cue-seeking task, subjects were returned to the operant chamber for 30 minutes where active lever pressing resulted in the illumination of the previously drug-paired cue light without a cocaine infusion.

### 16S sequencing & SCFA Metabolomics

Thirty minutes following completion of the cue-induced cocaine-seeking task, rats were sacrificed by rapid decapitation, and cecal contents were removed and flash frozen on dry ice. Samples were processed for 16S sequencing and SCFA metabolomics as described previously with full details in **Supplemental Materials and Methods**.

### RNA-sequencing analyses

Rats were euthanized as above, and the nucleus accumbens core was rapidly dissected on ice using the anterior commissure as an anatomical guide. RNA was isolated and cDNA libraries were prepared according to standard protocols. Libraries were sequenced with 150 nucleotide paired-end reads. Filtered reads were mapped to the mouse genome using STAR, and differential gene expression was performed using the DeSeq2 package with significantly regulated genes identified with the predetermined criteria of an FDR adjusted *p* value <0.2. Full methodological details in **Supplemental Materials and Methods**.

### Statistical Analysis / Figures

For analysis of self-administration acquisition and within-subject dose-response, repeated measures ANOVA or mixed effects analyses were utilized as appropriate. For PICRUSt, abundances were z-scored and analyzed using an ANOVA with a Tukey post-hoc. For pairwise behavioral comparisons, two-tailed Student’s t-tests were used. Analysis of RNA-sequencing and pathway data are as above. Figures were created using BioRender.com with permission to publish.

## Results

### Experiment 1: Microbiome depletion increases responding for low dose cocaine

To assess the effects of microbiome manipulation on drug self-administration, our previously established protocol of control (**H_2_O**) or non-aborsborable antibiotic-treated (**Abx**) animals (**Fig. 1A**) was used. Following the initial treatment period, rats were trained to self-administer cocaine on a fixed ratio 1 (FR1) reinforcement schedule at a dose of 0.8mg/kg/infusion. Initial studies found that microbiome manipulation affected the development of conditioned place preference and locomotor sensitization to low but not high dose cocaine^11^, so this higher training dose was selected to allow for equal acquisition of the response and to allow for subsequent testing of drug-seeking at different doses.

**Fig. 1.**
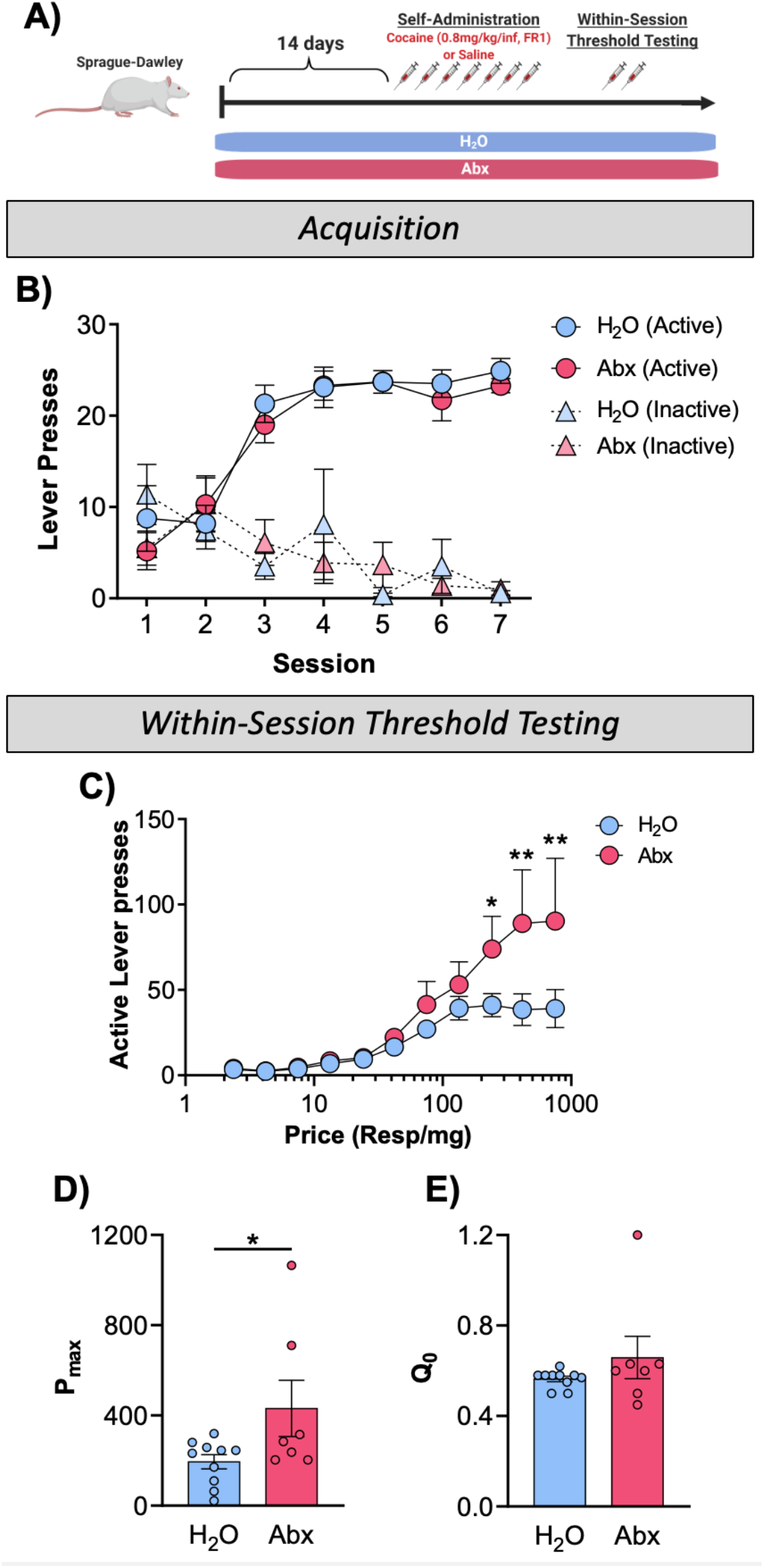
Abx effects on cocaine taking behaviors. **(A)** Schematic diagram of experimental procedures. **(B)** Acquisition of FR1 cocaine self-administration. Both H_2_O and Abx groups demonstrated robust and equal acquisition of active lever pressing (Main effect of day: *p* < 0.0001). **(C)** Dose-response curve for within-session threshold task demonstrated significant price x treatment interaction (*p* = 0.006) with Abx rats pressing more for lower doses. **(D)** Calculation of maximal effort animals will exert for cocaine, as measured by the P_max_ calculation, was increased in Abx rats (*p* = 0.048). **(E)** Drug intake at minimally constraining price (Q_0_) was not significantly different between groups. All data presented as mean ± SEM; * *p* < 0.05; ** *p* < 0.01.

Both the H_2_O and Abx groups acquired self-administration with a significant main effect of day (**Fig. 1B**; F_(6,87)_=30.79; *p*<0.0001), but no effect of treatment (*F*_(1,15)_=0.36 *p*=0.56) or any interaction (F_(6,87)_=0.53; *p*=0.79). Similarly, there were no treatment effects for inactive lever pressing (*F*_(1,17)_=0.009, *p*=0.93). Building from this foundation of equal administration between groups, animals were then tested on the within-session threshold task^42^ to assess motivation for cocaine^43,44^. **Figure 1C** shows the dose-response curve in which the cocaine dose delivered per lever press decreases over the session – on the x-axis increases in “price” are the number of presses required per mg of cocaine. There was a significant effect of price within these sessions, as was expected (F_(10,150)_=16.91; *p*=0.006). There was no significant main effect of antibiotic treatment (F_(1,15)_=2.61; *p*=0.13), but there was a significant price x treatment interaction owing to the fact that Abx-treated animals showed more robust responding for higher prices (i.e. lower dose/infusion) of cocaine (F_(10,150)_=2.596;*p*=0.006).

The power of the threshold task is that it assesses motivation and dose-response intake within a single session. Animals will maintain a preferred level of drug when the cost of drug intake is low. As price increases, animals are unable to maintain this dose and reduce their intake^42,43^. The inflection point at which this happens is P_max_, the maximal price that the animal is willing to pay to maintain a chosen concentration of drug. Abx-treated animals exhibited a significant increase in P_max_ (**Fig. 1D;** t=2.153, *p*=0.048). We also measured drug intake at a minimally constraining price, *Q_0_* (**Fig. 1E;***p*=0.25). There was no significant change, likely owing to similar drug intake between groups at higher doses when low effort was required to maintain intake.

### Experiment 2: Microbiome depletion increases cue-induced drug-seeking after abstinence

We examined whether antibiotic treatment would affect cocaine-seeking behavior in a seeking model of cocaine use disorder. A new cohort of rats was split into two groups, receiving either control (**H_2_O**) or antibiotics (**Abx**) for the duration of the study (**Fig. 2A**). For these experiments animals were trained to self-administer cocaine for 2 weeks. They Next underwent 21 days of home cage abstinence followed by a cue-induced cocaine-seeking task. Consistent with findings from Experiment 1, no group differences were observed in active (**Fig. 2B**; *F*_(1,11)_=3.39, *p*=0.09) or inactive (**Fig. 2B**; *F*_(1,11)_=0.28, *p*=0.61) lever pressing between subjects that received either antibiotics or control. A main effect of time was observed, indicating that both groups acquired cocaine self-administration (**Fig. 2B** active lever *F*_(3,33)_=6.98, *p*<0.001). No significant time by treatment interaction was observed (**Fig. 2B** active lever *F*(_13,138_)=0.42, *p*=0.96; **Fig. 2B** inactive lever; *F*_(13,137)_=0.92, *p*=0.53).

**Fig. 2.**
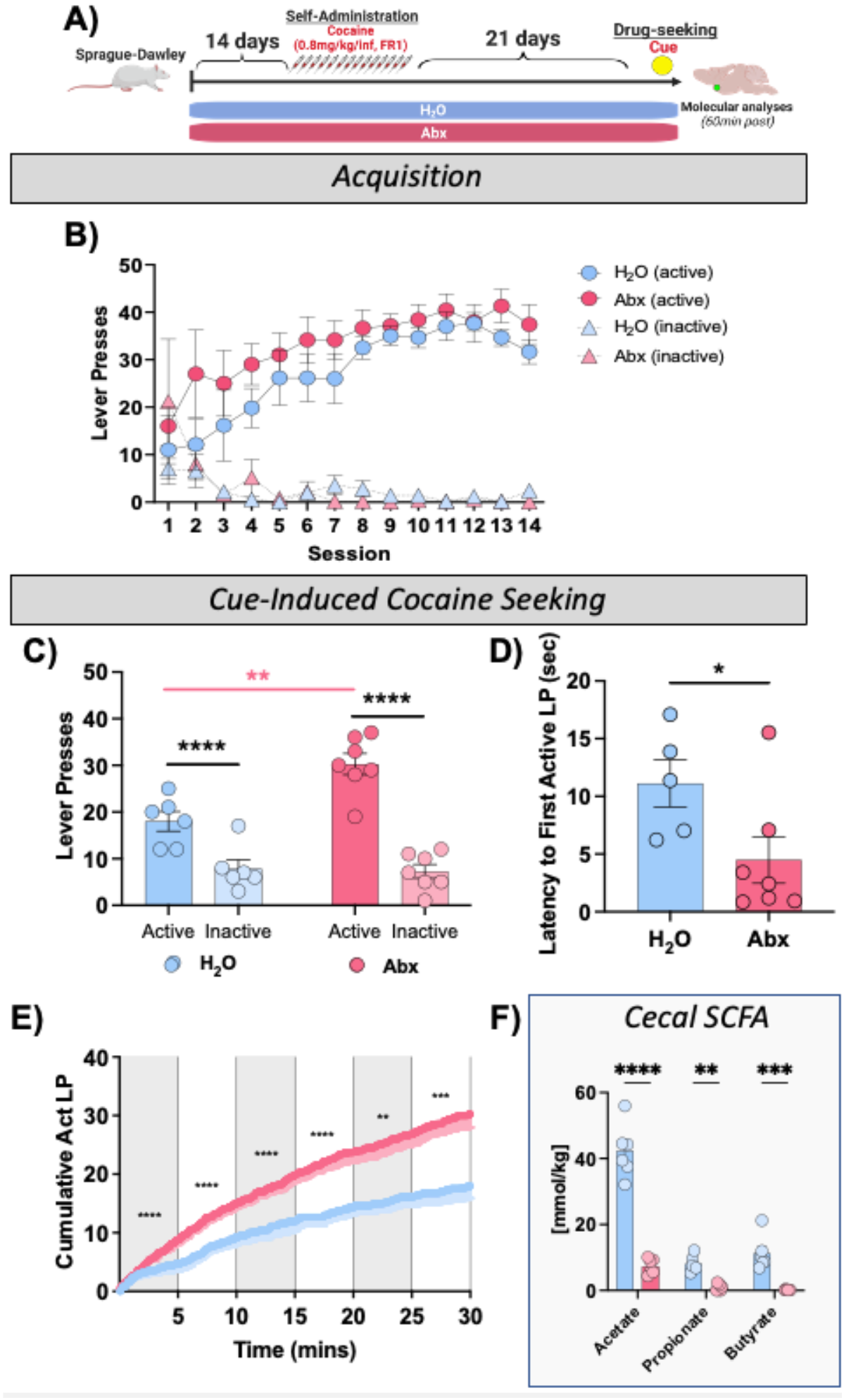
Abx increases on cocaine-seeking after abstinence. Schematic diagram of experimental procedure for this experiment. **(B)** Acquisition of FR1 self-administration. Both H_2_O and Abx groups showed robust acquisition of active lever pressing (Main effect of day: *p* < 0.0001). **(C)** On a cue-induced cocaine seeking task, both treatment groups showed robust lever pressing on the previously active lever – but pressing on the previously active lever was markedly increased in Abx-treated rats (red asterisks, p = 0.001). **(D)** The latency to first previously active lever press was significantly lower in Abx-treated rats (p = 0.049). **(E)** Analysis of cumulative active lever pressing across the session in five-minute bins shows Abx-treated rats had higher lever pressing in each bin measured. **(F)** Antibiotic treatment led to robust depletion of all three major SCFA in cecum. All data presented as mean ± SEM; * *p* < 0.05; ** *p* < 0.01; *** *p* < 0.001; **** *p* < 0.0001

On the cue induced seeking task, while both groups showed robust preference for the previously active lever (**Fig. 2C;** F_(1,22)_=69.60; *p*<0.0001), there was a main effect of antibiotics (F_(1,22)_=8.72; *p*=0.007), and a treatment x lever interaction (F(_1,22_)=10.42; *p*=0.004). Tukey’s post-hoc analysis demonstrated that this interaction was due to Abx-treated subjects pressing at much higher levels for the previously active lever (**Fig. 2C Red asterisks;***p*≤0.001) compared to H_2_O-treated controls, with no treatment effect on the inactive lever (**Fig. 2C;***p*=0.99). Post-hoc analysis further demonstrated that both Abx-treated (**Fig. 2C;***p*<0.0001) and control subjects (**Fig. 2C;***p*=0.012) exhibited increased responding for the previously active lever compared to the inactive lever, as expected.

As an additional measure of drug-seeking, we analyzed the latency to first lever press in response to a return to the chamber. Abx-treated subjects showed decreased latency to their first active lever press relative to H_2_O-treated controls (**Fig. 2D;***t*(_10_)=2.24, *p*=0.049). Examination of responding over the duration of the session was performed by analysis of cumulative responding for the previously active lever in five-minute bins for the duration of the 30 minute session. Abx-treated rats exhibited significantly increased sampling of the active lever at all time bins throughout the duration of the session (**Fig. 2E**; *p*<0.01 for all bins).

To assess for underlying molecular effects of microbiome depletion, we also quantified the three most abundant short-chain fatty acids (**SCFA**). We found that as expected antibiotic treatment robustly reduced levels of all three SCFA tested (**Fig. 2F**; Effect of treatment: F_(1,30)_=161.3, *p*<0.0001; Effect of analyte: F_(2,30)_=88.87, *p*<0.0001; Interaction: F_(2,30)_=38.87, *p*<0.0001).

### Experiment 3: SCFA repletion reverses the effects of antibiotics on cocaine-seeking after abstinence

Given that SCFA were decreased following antibiotic treatment and that prior studies demonstrate SCFA repletion reverses effects of antibiotic treatment on mouse conditioned place preference for cocaine and morphine^11,16^, we hypothesized that SCFA repletion would likewise reverse the effects of antibiotic treatment on cocaine-seeking. The experimental design was similar to Experiment 2 except subjects received either SCFA alone or a combination of SCFA+Abx in their drinking water **(Fig. 3A)**.

**Fig. 3.**
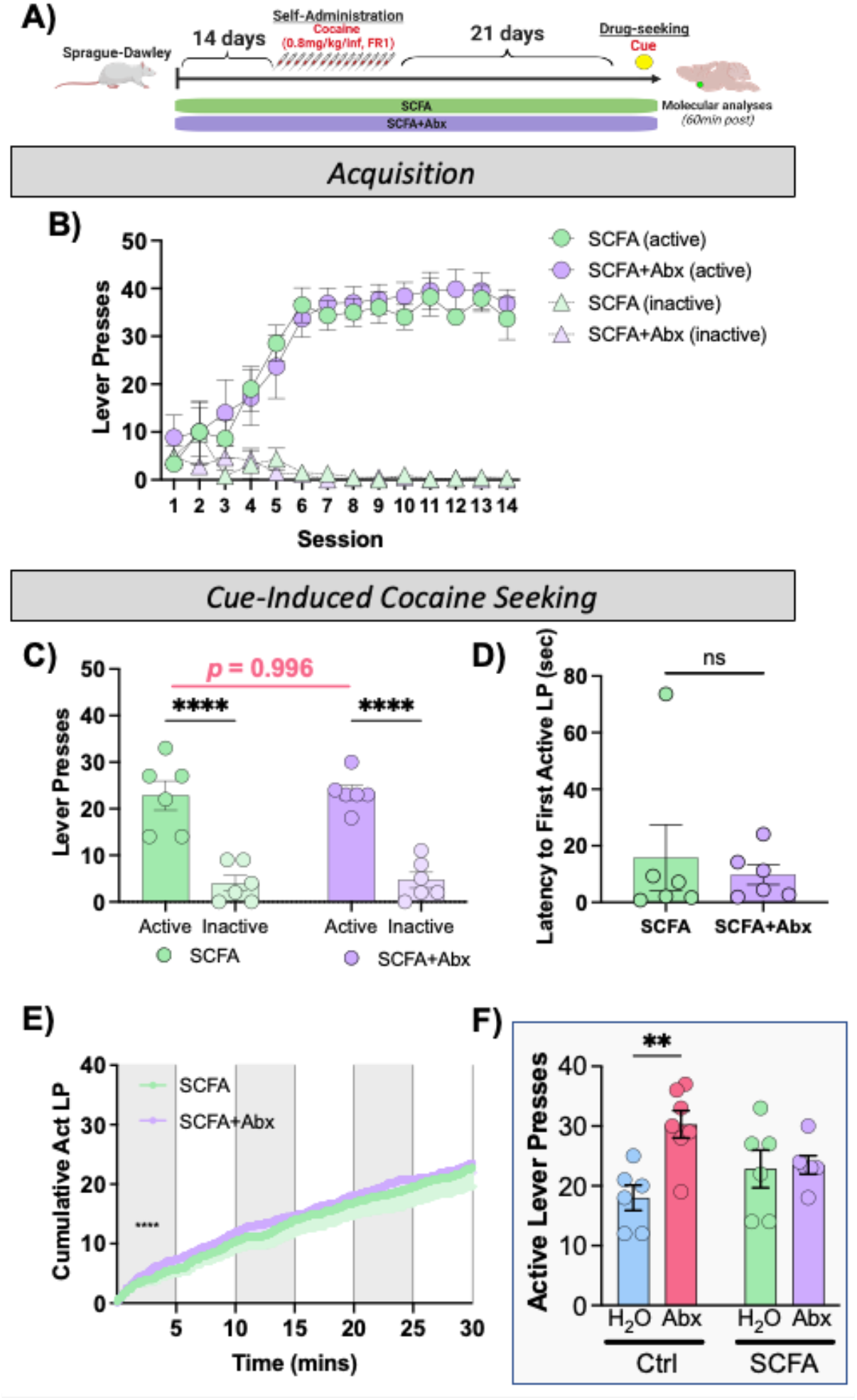
SCFA repletion reverses effects of microbiome depletion on drug-seeking. **(A)** Schematic diagram of experimental procedure. **(B)** SCFA and SCFA+Abx rats both acquires FR1 cocaine self-administration (Main effect of time: *p* < 0.0001). **(C)** On a cue-induced seeking task both groups showed robust responding on the previously active lever. However, in this case, rats treated with SCFA+Abx show equal total active responding to SCFA controls. **(D)** SCFA+Abx rats showed a trend toward increased latency to first response, but the effect was not significant. **(E)** Cumulative lever pressing over time shows parallel trajectories of responding between the two groups aside from the first five minutes. All data presented as mean ± SEM; **** *p* < 0.0001

Again, during acquisition no group differences were observed in either active (**Fig. 3B**; *F*_(1,11)_=0.29, *p*=0.60) or inactive (**Fig. 3B**; *F*_(1,11)_=0.02, *p*=0.89) lever pressing between subjects that received either SCFA or SCFA+Abx. Both groups acquired cocaine self-administration, as indicated by a main effect of time (**Fig. 3B** active *F*_(2.6,27.6)_=21.03, p<0.0001; inactive *F*_(2.7,28.8)_=3.09, *p*=0.048). Time by treatment interaction was not observed (**Fig. 3B** active *F*_(13,139)_=0.32, *p*=0.98; inactive *F*_(13,139)_=0.83, *p*=0.63).

After 21 days of abstinence, both groups exhibited cocaine-seeking with a main effect of active lever pressing (**Fig. 3C**; *F*_(1,22)_=72.63, *p*<0.0001). As hypothesized, SCFA repletion attenuated the effects of antibiotics on cue-induced cocaine-seeking. There was no main effect of antibiotic treatment (**Fig. 3C**; *F*_(1,22)_=0.85, *p*=0.37), nor was there a significant interaction (**Fig. 3C**; *F*_(1,22)_=0.027, *p*=0.87). Tukey’s post-hoc analysis demonstrated no differences in responding for either the previously active lever (**Fig. 3C**; *p*=0.87) or the inactive lever (**Fig. 3C**; *p*=0.95). Post-hoc analysis further demonstrated that both SCFA+Abx-treated (**Fig. 3C**; *p*<0.0001) and SCFA-treated subjects (**Fig. 3C**; *p*<0.0001) exhibited increased responding for the previously active lever compared to the inactive lever, as expected.

No differences were observed in latency to first active lever press (**Fig. 3D**; *t*_(11)_=1.73, *p*=0.11). Cumulative responding analysis revealed that SCFA+Abx rats exhibited increased responding for the previously active lever only during the first five minutes of the session (**Fig. 3E;** F_(1,3896)_ =289.0, *p*<0.0001). No differences in responding were observed for the other time bins across the 30-minute session.

### Microbiome depletion decreases microbial diversity and predicted SCFA metabolism

To explore the effects of antibiotic-treatment and SCFA repletion microbial diversity, we performed 16S sequencing on cecal contents from subjects that underwent behavioral testing for cue-induced cocaine-seeking following abstinence (**Figs. 2A & 3A**). As expected, there was a robust main effect of antibiotics on alpha diversity (**Fig. 4A;** F_(1,20)_=3194, *p*<0.0001). Interestingly, there were main effects of SCFA treatment (F_(1,20)_=23.38, *p*=0.0001) and an antibiotics x SCFA interaction (F_(1,20)_=23.03, *p*=0.0001). On post-hoc testing there were between group differences for Abx-treated and non-Abx-treated groups. While there was no difference between control H_2_O-treated and SCFA-treated rats (*p*=0.98), there was a modest but significant difference between Abx-treated and Abx+SCFA-treated rats for alpha diversity (**Fig. 4A** red/purple comparison; *p*<0.0001).

**Fig. 4.**
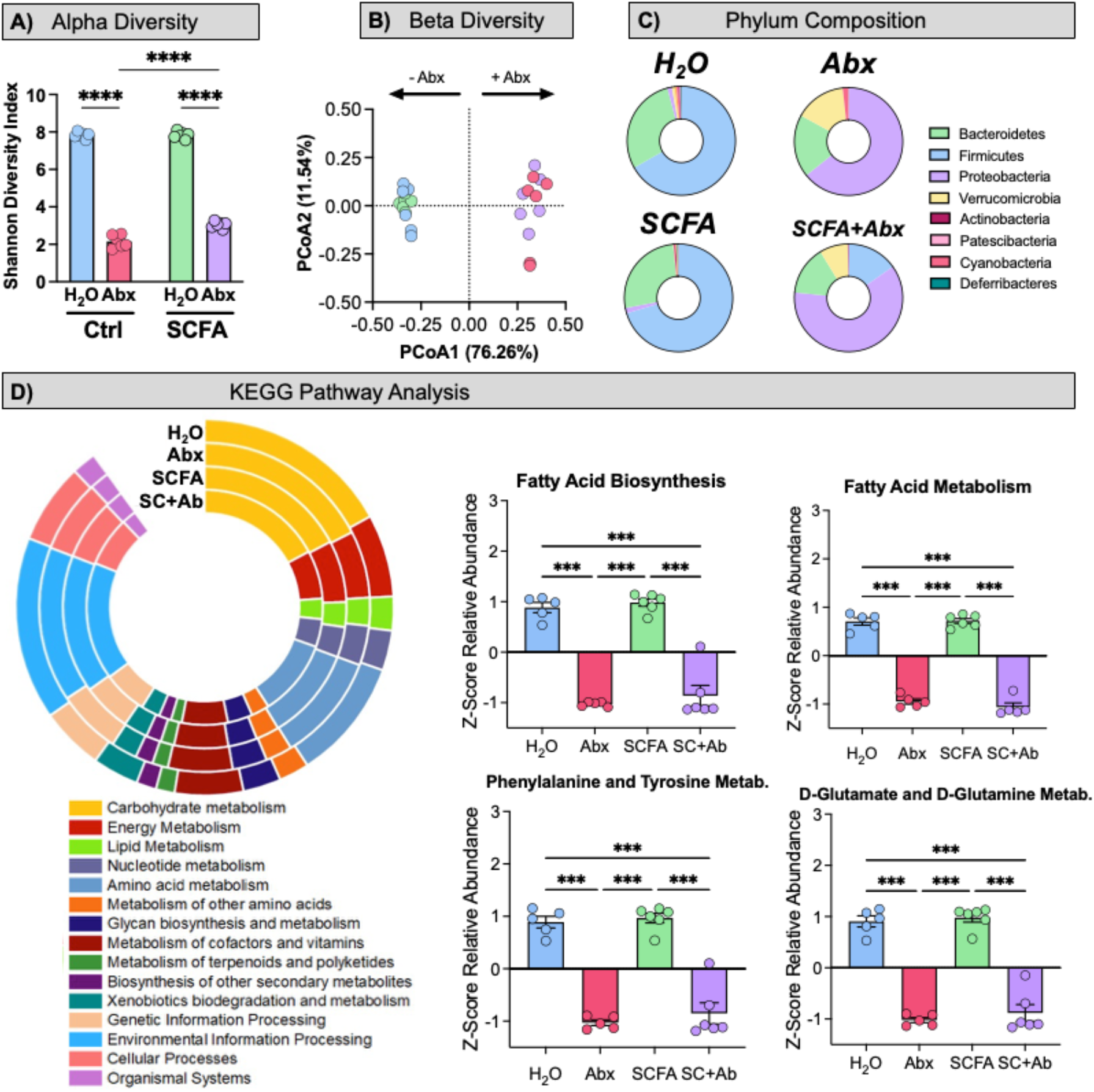
Effects of Abx and SCFA on microbiome composition and function. **(A)** Using the Shannon measure of alpha diversity, a measure of within sample diversity, there are marked reductions in diversity with antibiotictreatment in both groups (Main effect of Abx *p* < 0.0001), and also a less robust but statistically significant main effect of SCFA treatment (*p* = 0.0001). **(B)** Weighted UniFrac dissimilarity analysis of beta diversity demonstrates antibiotic treatment is the main driver of between subjects diversity changes. **(C)** Donut plots of relative phylum composition show unique changes in bacterial composition between groups. Importantly, these plots show percentage of total, not absolute quantification. **(D)** High-level overview of PICRUSt data highlights KEGG abundances between treatment groups. KEGG pathway relative abundance is represented in a chipped-donut plot with group labels at the top of the plot. Highly differential pathways involved in fatty acid biosynthesis, fatty acid metabolism, phenylalanine and tyrosine metabolism, and glutamate metabolism are shown as z-scores ± SEM to the right. All plots demonstrate a down regulation in the Abx and Abx+SCFA groups compared to H_2_O and SCFA groups.

Beta diversity analysis using a weighted UniFrac dissimilarity matrix showed that the primary driver of diversity was antibiotic treatment (**Fig. 4B** x-axis accounts for 76.26% of variability); within these groups the control and SCFA treatment groups largely co-clustered. Antibiotic treatment resulted in significant decreases in the relative abundance of Actinobacteria, Deferribacteres, Firmicutes, Patescibacteria and Tenericutes, and additional unclassified phyla (**Fig. 4C**). SCFA treatment groups looked similar to their control counterparts, but with SCFA+Abx displaying some shifts in Proteobacteria, Bacteroidetes, and Firmicutes relative to Abx-treated subjects.

To gain more granular insight into the relative changes induced by antibiotics and SCFA treatment, we performed PICRUSt2 analysis. PICRUSt2 predicts the functional potential of bacterial communities based on sequencing profiles. These predictions are supported by integration into the Kyoto Encyclopedia of Genes and Genomes (**KEGG**) orthologs (**KO**) and Enzyme Commission numbers (**EC**) to explore related functional changes. A high-level overview of abundance of KEGGs categorized by metabolic function is shown in **Fig. 4D - left** as a chipped donut plot of KEGG abundance categorized by KEGG mapper. To further explore relevant pathways, we evaluated KOs related to fatty acid biosynthesis and metabolism as well as essential amino acids required for neurotransmitter production. Significant differences in abundance following antibiotic treatment were observed in the KOs for Fatty Acid Biosynthesis (**Fig. 4D-Top Middle;** F_(3,18)_=73.86; *p*<0.001), Fatty Acid Metabolism (**Fig. 4D – Top Right;** F_(3,17)_=218.8; *p*<0.001), Phenylalanine, Tyrosine, and Tryptophan Metabolism (**Fig. 4D - Bottom Middle** F(_3,18_)=64.18; *p*<0.001), and D-Glutamate and D-Glutamine Metabolism (**Fig. 4D-Bottom Right;** F_(3,18)_=91.43; *p*<0.001), respectively. Post-hoc analysis for each of the aforementioned KOs revealed significantly decreased biosynthesis and metabolism in the Abx and SCFA+Abx groups compared to the H_2_O and SCFA groups. Further analysis of microbiome data is available as **Supplemental Figures 2–4**.

### Microbiome depletion alters gene expression in the nucleus accumbens

Although many brain regions contribute to drug reward and the development of substance use disorders, the nucleus accumbens (**NAc**) is the most heavily implicated as a primary reward center in the brain, playing key roles in drug-seeking after abstinence^45,46^. Previous work demonstrates that manipulations of the microbiome can markedly affect transcriptional regulation in the brain^11,16,31,47,48^. To explore the impact of microbiome depletion and SCFA repletion on gene expression in brain reward circuitry, we performed full transcriptomic RNA sequencing on NAc punches from subjects that underwent behavioral testing for cue-induced cocaine-seeking following abstinence (**Fig. 2A & 3A**). When compared to the H_2_O controls, we find that all three treatment groups exhibited altered gene expression with 551 DEGs in the Abx group, 262 in the SCFA+Abx group, and 149 in the SCFA group (**Fig. 5A**). Significant gene ontologies of interest from each group are provided as **Supplemental Fig. 5**; full list of significant GO terms from each is available as **Supplemental Tables 1-3**. To dissect the individual differences and overlap between the treatment groups, we created an UpSet plot^49^ (**Fig. 5B**). The number of statistically significant genes in each group is shown on the lower left horizontal bar graph. Intersections between groups are demonstrated in the right side of the graphic with filled in circles in the dot plot representing the genes found in each individual group or combination of groups. This analysis demonstrates that Abx-treated mice have the highest number of differentially expressed genes relative to controls (**Fig. 5 - right**) – and importantly that the majority of these significant genes are unique to the Abx-treatment group (**Fig. 5B - right, red bar, 421 for Abx)**. Animals treated with SCFA+Abx have a significant difference of only 123 unique genes, suggesting that addition of the SCFA to animals with depleted microbiomes led to normalization of transcriptional regulation as well as drug-seeking behaviors.

**Figure 5.**
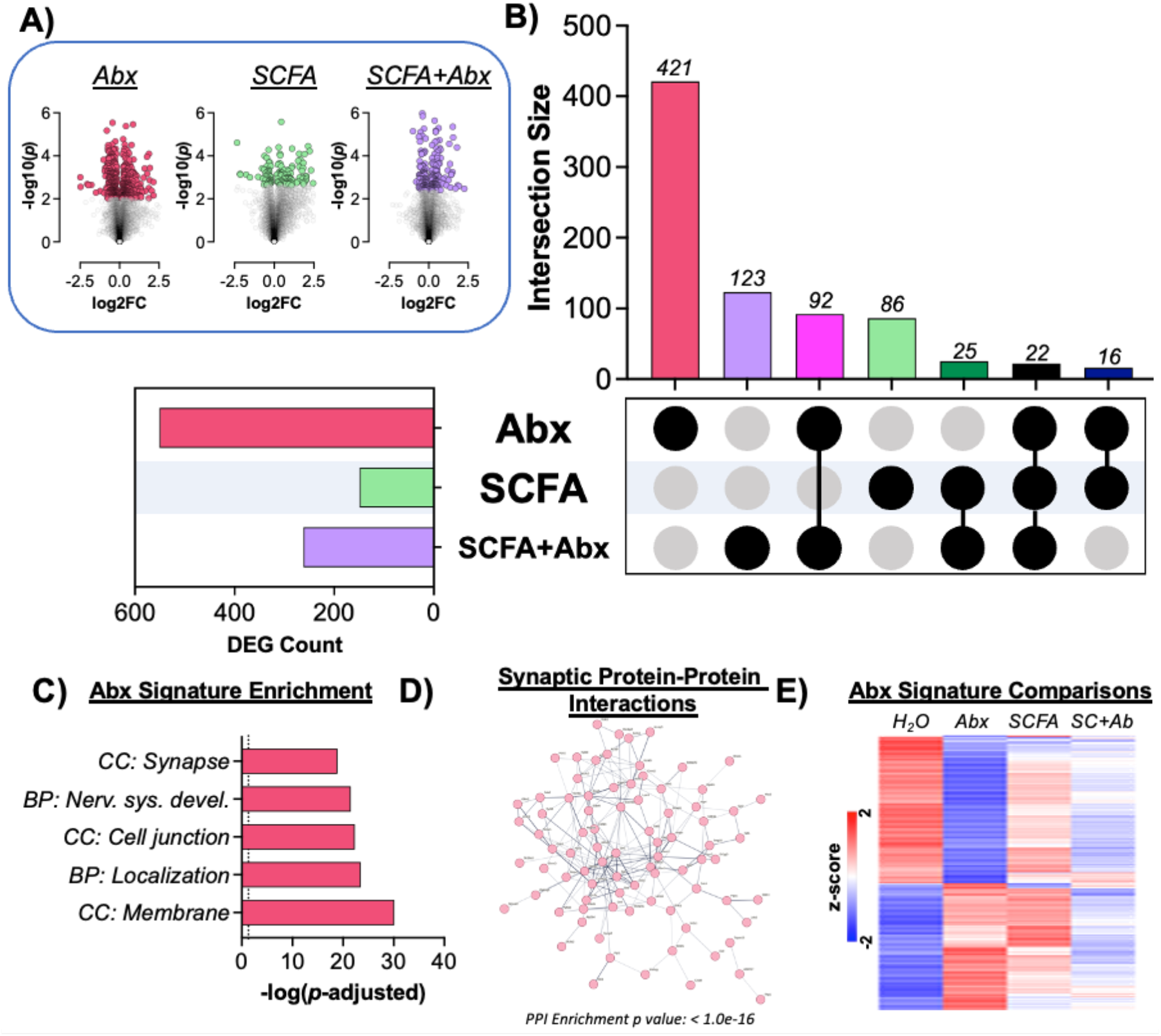
RNA sequencing analysis of NAc after drug seeking. **(A)** Volcano plots of gene expression in all three treatment groups relative to H_2_O controls. Colored points in each panel are statistically significant after FDR correction. **(B)** UpSet plot of unique and overlapping gene expression patterns between each group. Left bar graphrepresents the number of significant genes in each treatment category. Right graph shows the number of genes in each treatment or treatment combination as indicated by filled circles. **(C)** Significant GO terms of interest from the unique “Abx Signature” gene list. **(D)** Predicted protein-protein interactions among genes from synapse and cell junctions gene lists. **(E)** Heatmap showing z-scored FPKM values of genes from the Abx signature list demonstrating moderation of SCFA+Abx compared to Abx only.

A detailed analysis of these “Abx Signature” genes that were uniquely altered in Abx-treated rats revealed marked alterations in genes related to synaptic pathways, nervous system development, and membrane function (**Fig. 5C** & **Supp. Table 4**). STRING analysis of predicted protein-protein interactions amongst these gene products in the synapse and cell junction pathways predicted changes in a highly interactive group of pre- and post-synaptic functions (**Fig. 5D**). Finally, heatmap analysis of the genes from this antibiotic signature shows that while these genes were all markedly different from H_2_O controls, there was a tempering effect particularly noted in the Abx+SCFA group (**Fig. 5E** - right column). These data demonstrate that depletion of the microbiome leads to marked transcriptional and behavioral changes – and that SCFA metabolite repletion can attenuate some of the molecular and behavioral phenotypes.

## Discussion

We show that antibiotic induced microbiome depletion has marked effects on both active cocaine self-administration and drug-seeking after abstinence. Rats treated with antibiotics show normal acquisition of cocaine self-administration for high dose (0.8mg/kg/inf) cocaine on an FR1 schedule. However, when using a within-session threshold task, we found that microbiome depletion was associated with enhanced drug intake at the lower bounds of the dose range. This suggests enhanced motivational properties of cocaine or a shift in the dose-response curve induced by microbiome depletion. While within-session threshold testing is a useful measure of drug-seeking during active use, we wanted to examine the role of gut-brain signaling in drug-seeking following abstinence using a relapse model. We find that rats with a depleted microbiome exhibit increased drug-seeking in response to a drug-paired cue following three weeks of forced abstinence. This is in line with the threshold task data, suggesting that depletion of the microbiome enhances the motivational properties of cocaine. Importantly, the effects of microbiome depletion could be reversed by repletion of SCFAs, suggesting a possible mechanism for the observed behavioral effects. When we examined how microbiome depletion and SCFA repletion affected transcriptional regulation following drug-seeking, we found that alterations in the microbiome and its metabolites played an important role in shifting the transcriptional landscape. Taken together, these findings demonstrate a clear role for the microbiome and its metabolites in drug-taking and seeking, laying the foundation for future translational work in this space.

Several years ago it was demonstrated that depletion of the gut microbiome led to enhanced cocaine conditioned place preference and locomotor sensitization at low – but not high – doses of cocaine^11^. These behavioral effects were also reversed by repletion of SCFA in antibiotic-treated mice. Lee and colleagues demonstrated that a shorter treatment with a different antibiotic cocktail led to a reduction in cocaine place preference at an intermediate dose^50^. Recent studies demonstrated production of gut microbiome-derived glycine can have marked effects on behavioral responses to cocaine^19^. Interesting recent work has even identified interactions between the microbiome and social stimuli in regulating cocaine place preference^18^. Studies of microbiome effects on non-stimulant drugs of abuse have also shown marked effects of the microbiome on behavior. Recently, we found that depletion of the microbiome using the same antibiotic regimen led to markedly decreased preference for morphine across a wide dose range, and that this microbiome effect was again reversable by SCFA repletion^16^. These results are interesting, as they use identical manipulations of the microbiome and its byproducts – but show opposite effects with opioids. A number of well-crafted studies have demonstrated that a diverse microbiome is important for normal development of tolerance to morphine over time; both germ-free and antibiotic-treated mice show markedly reduced tolerance for morphine^21,32^. All of these previous studies have utilized shorter term experimenter-administered drugs. The studies in this manuscript are the first to utilize voluntary cocaine self-administration and to employ a method for examining drug-seeking in a model for relapse.

One potential mechanism by which manipulations of the gut microbiome can alter neurobiological plasticity and subsequent behavior is via effects on transcriptional regulation in the brain. Previous studies have demonstrated that lack of a diverse gut microbiome alters transcriptional regulation in the amygdala and prefrontal cortex following fear conditioning^47,48,51,52^. Microbiome depletion significantly alters regulation of neuroplasticity-related genes in the NAc following cocaine exposure^11^. Likewise, in opioid-exposed animals, microbiome depletion altered immediate early gene activation patterns^53^ in addition to transcriptome-wide dysregulation^16^. Experiments presented here have the important distinction that transcriptional profiling was performed three weeks after final drug exposure and following a cue-induced drug-seeking task. As a likely result of this difference, transcriptional effects are more modest than those seen immediately after drug intake (**Fig. 5**). However, these findings are similar to those seen after a fear conditioning task in which there was a significant microbiome x behavior interaction in transcriptional regulation^48^.

Importantly, repletion of the microbiome-derived SCFA metabolites leads to reversal of the microbiome effects on drug-seeking and alters transcriptional regulation in the NAc. We examined these metabolites as there is a robust literature demonstrating SCFA as key gut-brain signaling molecules^54^. These molecules, which are produced by gut bacteria in the process of fermentation of dietary fiber and are nearly completely eliminated by antibiotic treatment (**Fig. 2F**), are implicated in myriad gut-brain signaling pathways from control of blood-brain barrier integrity and microglial function to regulation of food intake^27,29,54,55^. Previously, SCFA are shown to regulate behavioral responses to both experimenter-administered cocaine and opioids^11,16^. There is robust evidence that the SCFA modulate transcriptional regulation in the brain. All of these small molecules function as histone deacetylase inhibitors, with butyrate and acetate having the strongest effects^56^. Recently gut-derived acetate was found to alter patterns of histone acetylation in the brain, affecting consolidation of memory and behavioral response to alcohol^57,58^. All SCFA regulate activation of the CREB transcription factor and alter expression of numerous gene products including tyrosine hydroxylase, c-Fos and enkephalin^59–65^.

Our understanding of how the gut microbiome and its resultant byproducts can alter brain and behavior is advancing at a rapid rate. These findings provide the first critical evidence that manipulation of the gut microbiome and its metabolites can alter cocaine self-administration and drug-seeking after relapse. These studies provide important foundational data to move gut-brain signaling towards further translational studies, which should examine how specific microbial composition and metabolite levels work to drive both drug-seeking and other motivated behaviors. Ultimately, there is strong potential for these microbial signaling pathways to be explored as either biomarkers or treatments for patients with debilitating substance use disorders. While much remains to be done, this work suggests strong potential for these pathways to be harnessed from the bench to the bedside.

## Acknowledgements

The authors would like to thank NIDA Drug Supply for provision of cocaine hydrochloride. We would like to thank Alexia Seba-Robles for assistance with animal care. Figures made with BioRender with full permission to publish.

## Author Contributions

D.D.K., E.S.C, and K.R.M. designed the experiments. K.R.M,, A.G., E.G.P, E.S.C., R.S.H., and D.D.K. performed experiments. K.R.M., S.S.S., A.G., O.G., E.S.C., R.S.H. & D.D.K. analyzed data. K.R.M. & D.D.K. wrote the manuscript. All authors provided critical edits and feedback of the finalized manuscript.

## Supplemental Methods

### Jugular Catheterization

Subjects were first anesthetized with a cocktail of ketamine 100mg/kg and xylazine 10mg/kg followed by placement of an indwelling catheter (Plastics One, Torrington, CT) into the right jugular vein. For all experiments, subjects were allowed to recover for 2-4 days before the start of cocaine self-administration. Catheters were flushed daily with heparinized saline to maintain patency. To avoid confound of additional antibiotic treatment, subjects did not receive post-catheterization IV antibiotics. No signs of surgical site infection were observed in any animals.

### Experiments 1-3: Cocaine Self-Administration: Acquisition

Cocaine self-administration was performed in standard operant conditioning chambers (MED Associates, St. Albans, VT) with single speed syringe pumps for drug delivery, two retractable levers, and two lights located above each lever. Subjects were food restricted starting one day prior to self-administration training and were continued on food restriction throughout the duration of self-administration training with 18g food/subject delivered once daily following session completion. Sessions were once daily and lasted for 3 hours. Sessions were initiated by the extension of the active and inactive levers. Subjects were trained to respond for cocaine on the “active” lever on a fixed ratio 1 schedule of reinforcement where one lever press resulted in the delivery of a 5.9s infusion of 0.8 mg/kg/infusion (0.1 ml) cocaine in sterile saline paired with concurrent illumination of the cue light located above the active lever. Each infusion was followed by a 20 second time out. Responses on the inactive lever were recorded, but had no programmed consequence. Inactive lever position was counterbalanced across groups.

### Experiment 1: Within Session Threshold Testing

Following acquisition of FR1 administration, a within session threshold test was performed to assess differences in cocaine motivation and consumption between groups. In this behavioral economics task, rats are given continuous access to cocaine while increasing the effort requirement to obtain the same amount of drug^1,2^. For this, subjects were maintained respond on an FR1 schedule for a descending series of 11 unit doses of cocaine, (421, 237, 133, 75, 41, 24, 13, 7.5, 4.1, 2.4, and 1.3 μg/infusion) with no timeout period following infusions or between 10 minute bins. Each dose of cocaine is available for 10 minutes, with each dose bin presented consecutively across the 110-min session. Responding during the first bin of the procedure is considered to reflect a loading phase and is not included in the analyses. Cocaine consumption as a function of price (number of active lever presses to obtain 1 mg cocaine) is plotted as a within-session dose response curve (**Fig. 1C**). The dose is high during the initial bins of the procedure, allowing the subject to consume a preferred level of cocaine with minimal effort. As the session continues and the dose is lowered across bins, the subject must exert increasing effort to maintain consistent drug intake.

Shifts in responding across the dose curve can be analyzed using behavioral economics principles, to assess a variety of behavioral economic measures as described previously^1,2^. Behavioral economic analysis was used to determine the parameters of maximal price paid (P_max_) and consumption at a minimally constraining price (Q_0_), as described previously^3–5^. Demand curves were generated by curve-fitting individual animals’ intake using an equation: log(Q) = log(Q_0_) + *k* × (*e*–*α* × Q_0_ × *C*–1)^6,7^. Previous work has demonstrated that P_max_ is correlated with break points on a progressive ratio schedule of reinforcement, confirming that the threshold procedure accurately assesses reinforcing efficacy^2,5^. The value *k* was set to 2 for all animals^6,7^.

#### Q_0_

*Q*_0_ is a measure of the animals’ preferred level of consumption. This can be measured when cocaine is available at low effort, or a minimally constraining price. This preferred level of consumption is established in the early bins of the threshold procedure.

#### P_max_

Price is expressed as the responses emitted to obtain 1 mg of cocaine, thus as the effort requirement is increased, the relative price to obtain cocaine also increases. As the session progresses, animals must increase responding on the active lever in order to maintain stable intake. P_max_ is the price at which the animal no longer emits enough responses to maintain intake. Thus, animals with higher P_max_ will increase responding to maintain cocaine levels farther into the demand curve; in other words, they will pay a higher price for cocaine.

### 16S Sequencing of the Microbiome

Microbial DNA was isolated from frozen cecal contents using the DNAeasy PowerSoil Kit (Qiagen 12888) per manufacturer’s protocol with the addition of a bead beating step to ensure complete and uniform lysis of bacterial cells. DNA concentration was measured via Nanodrop1000, and at least 400ng of DNA was sent to LC Sciences for 16S RNA sequencing. Sequencing was performed as previously described^8^. Briefly, sequences with ≥ 97% similarity were assigned to the same observed taxonomic units (OTU). OTUs were identified and taxonomy assigned by comparing representative genetic sequencing for each OTU to reference bacterial genomes from the Ribosome Database Project^9^. Quantitative Insights Into Microbial Ecology 2 (QIIME2) software was used to assess alpha and beta measures of microbial diversity^10^. OTU counts were used to generate Simpson and Shannon alpha diversity indices after rarefaction^10^. Principle component analysis plots were constructed based on Unifrac distance as a measure of beta diversity^10^. Phylogenetic Investigation of Communities by Reconstruction of Unobserved States (PICRUSt2) software was used to predict the functional profiles of microbial taxa based on observed OTUs^11,12^. For heatmaps and bar graphs PICRUSt2 outputs were quantified and then z-scored for quantitative comparisons.

### SCFA Metabolomics

To quantify cecal SCFA levels, rats were placed on Abx, SCFA, SCFA+Abx, or maintained on control H_2_O for 7 weeks prior to rapid decapitation and removal of cecal contents as described. SCFA were quantified using a Water Acquity uPLC System with a Photodiode Array Detector and an autosampler. Samples were analyzed on a HSS T3 1.8 μm 2.1×150 mm column with a flow rate of 0.25 mL/min, an injection volume of 5 uL, a column temperature of 40°C, a sample temperature of 4°C, and a run time of 25 min per sample. Eluent A was 100 mM sodium phosphate monobasic, pH 2.5; eluent B was methanol; the weak needle wash was 0.1% formic acid in water; the strong needle wash was 0.1% formic acid in acetonitrile, and the seal wash was 10% acetonitrile in water. The gradient was 100% eluent A for 5 min, gradient to 70% eluent B from 5-22 min, and then 100% eluent A for 3 min. The photodiode array was set to read absorbance at 215 nm with 4.8 nm resolution. Samples were quantified against standard curves of at least five points run in triplicate. Standard curves were run at the beginning and end of each metabolomics run. Quality control checks (blanks and standards) were run every eight samples. Concentrations in the samples were calculated as the measured concentration minus the internal standard; the range of detection was at least 1 – 100 μmol/g stool.

### RNA Processing & cDNA Library Preparation

RNA was isolated using RNeasy kits (Qiagen-#74106) with on-column DNAase digestion (#79254) per manufacturer’s protocol. The integrity and purity of total RNA were assessed using Agilent Bioanalyzer and OD_260/280_ using Nanodrop. cDNA libraries for full transcriptomic RNA-seq were prepared using the NEBNext Ultra II Directional RNA Library Prep Kit for Illumina (E7765). Starting with 1.0 ug total RNA per sample, mRNA was isolated using the NEBNext Poly(A) mRNA Magnetic Isolation Module (E7490) and fragmented. cDNA was synthesized using random primers, end-repaired, adaptor ligated, and purified. Samples were multiplexed using NEBNext Multiplex Oligos for Illumina Kit (E7335, E7500) with single index 6-bp barcodes introduced at the end of the adaptors during PCR amplification per manufacturer’s protocol. The libraries were sequenced by Novogene on the Illumina HiSeq system with 150 nucleotide paired end reads.

### RNA-Sequencing Data Preprocessing and Analysis

Raw reads were subjected to quality control and filtered to remove low quality reads (i.e. either (1) uncertain nucleotides constituting >10% of either read, (2) nucleotides with base quality <20 constituting >50% of the read, or (3) containing adaptor contamination). Filtered reads were mapped to the mouse reference genome using STAR^13^. Gene expression level was determined by counting the reads that map to exons or genes. Transcripts per million of transcript sequence per illion base pairs sequenced (TPM) was calculated to estimate gene expression levels. For pairwise comparisons, aligned reads were analyzed for differential gene expression using DeSeq2 analysis package via Network Analyst^14^. Significantly regulated genes were identified with the predetermined criteria of an FDR adjusted *p* value <0.2. Lists of significantly regulated genes were utilized for both G:Profiler^15^, and significantly regulated biological pathways were determined using a FDR correction threshold of *p* adj <0.05.

## Supplemental Results

MetaCyc^16^ is a pathway analysis database that can be used to predict changes in metabolic pathways from 16S sequencing data downstream of PICRUSt2^17^. This software operates with two types of pathways, base pathways that represent individual metabolic pathways, and super pathways which combine sets of the base pathways into larger metabolically related pathway. Fatty acid metabolism pathways were interrogated in three groups, acetate metabolism, butanoate metabolism, and propanoate metabolism. **Figure S2A-2D** are related to acetate metabolism, **2E-2H** are related to butanoate metabolism, and **2I-L** are related to propanoate metabolism. Globally, butanoate metabolism is down-regulated in Abx treated groups compared to H_2_O and SCFA treated groups. There appears to be some compensation as the Abx group has upregulated fatty acid super pathway (**2A**) abundance, as well as increased acetate production from hexitol fermentation (**2B**), but a decrease in pyruvate fermentation to acetone (**2C**), as well as acetate and lactate (**2D**). Propanoate metabolism is relatively unaffected, except for an increase in 3-phenylpropanoate degradation in the Abx group.

MetaCyc pathways are the highest-level output for PICRUSt and are generated by structured mapping of EC gene families to pathways. For **Figure S3**, a z-score of the relative abundance of EC values was taken to compare across groups, then EC values are mapped onto SCFA metabolism and biosynthesis pathways. The colors are relative to the Abx group as compared with the H_2_O control. If the Abx EC is upregulated, the EC is colored green. If the Abx EC is downregulated, the EC is colored red. If the Abx EC is not significantly different from the H_2_O group, the EC is colored gray. Intermediate metabolites are represented as blue ovals. A significant increase or decrease was considered if (*p*<0.05). Many ECs related to butyrate (butanoate) production either from succinate or pyruvate are downregulated compared to the H_2_O group suggesting depletion of the microbiome reduces functional efficacy of these pathways.

To provide a holistic view of all super pathways assessed by MetaCyc, we quantified relative abundances which were then z-scored and sorted with unsupervised hierarchical clustering in **Fig. S4**. As previous, the main effects are driven by antibiotic treatment. The top highly differentially abundant pathways involve caprolactam degradation, propanoate metabolism, butanoate metabolism, amino acid metabolism.

## Supplemental Figures

**Supplemental Figure 1.**
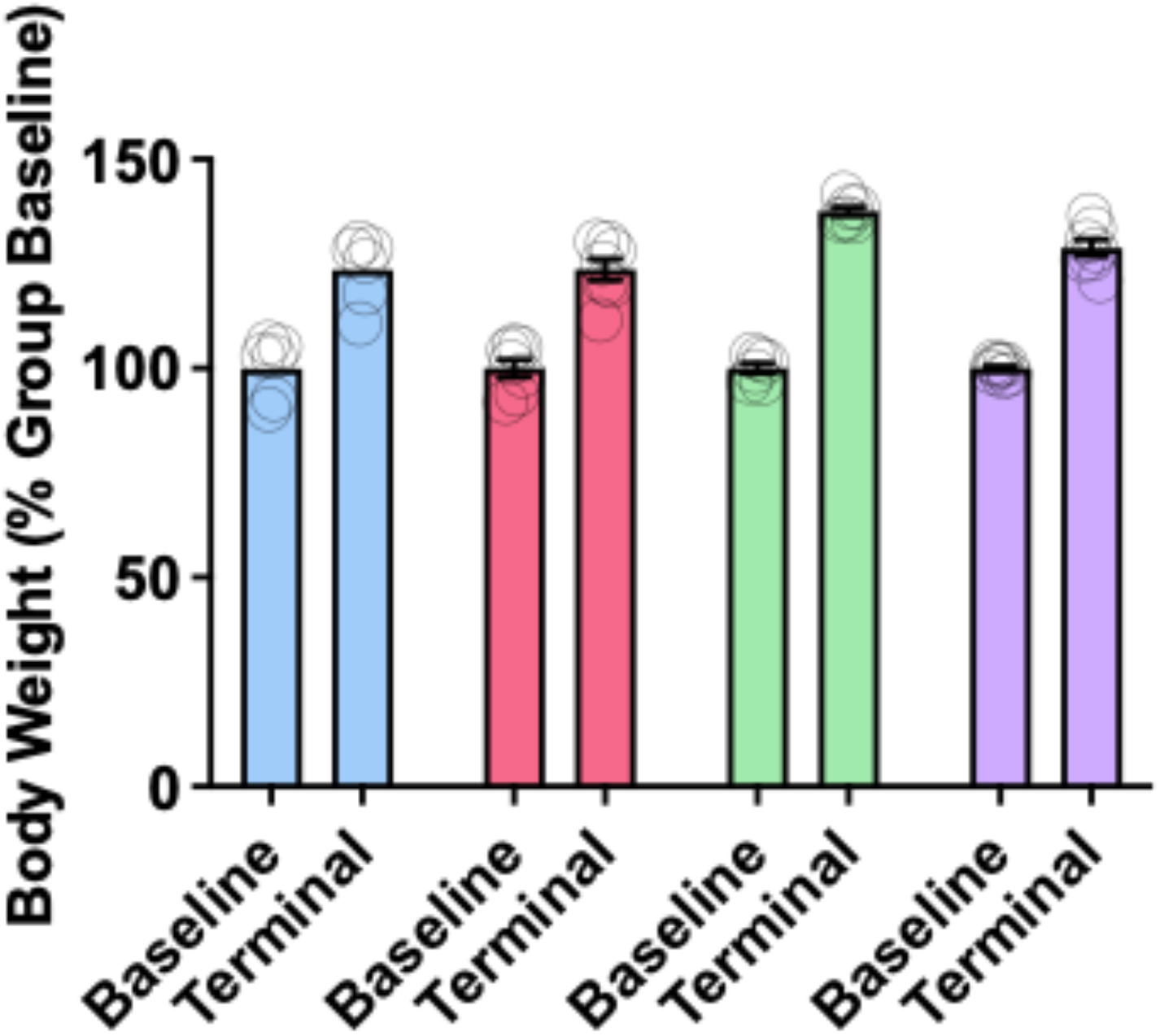
Abx and SCFA metabolites do not affect weight gain over time. To ensure that animals were taking in adequate fluid and maintaining healthy body composition over time, a subset of animals was weighed over the course of the experiments. Animals in all groups showed robust weight gain over the course of the experiment (Main effect of time: F(_1,22_) = 1631; *p* <0.0001), however there was no main effect of treatment (F_(3,22)_ = 2.7; *p* =0.07). There was a significant time x treatment interaction (F(_3,22_) = 20.95; *p* <0.0001), but no individual comparisons were significant on between group post-hoc testing (*p* ≥ 0.11 for all). N = 6-7/group.

**Supplemental Figure 2.**
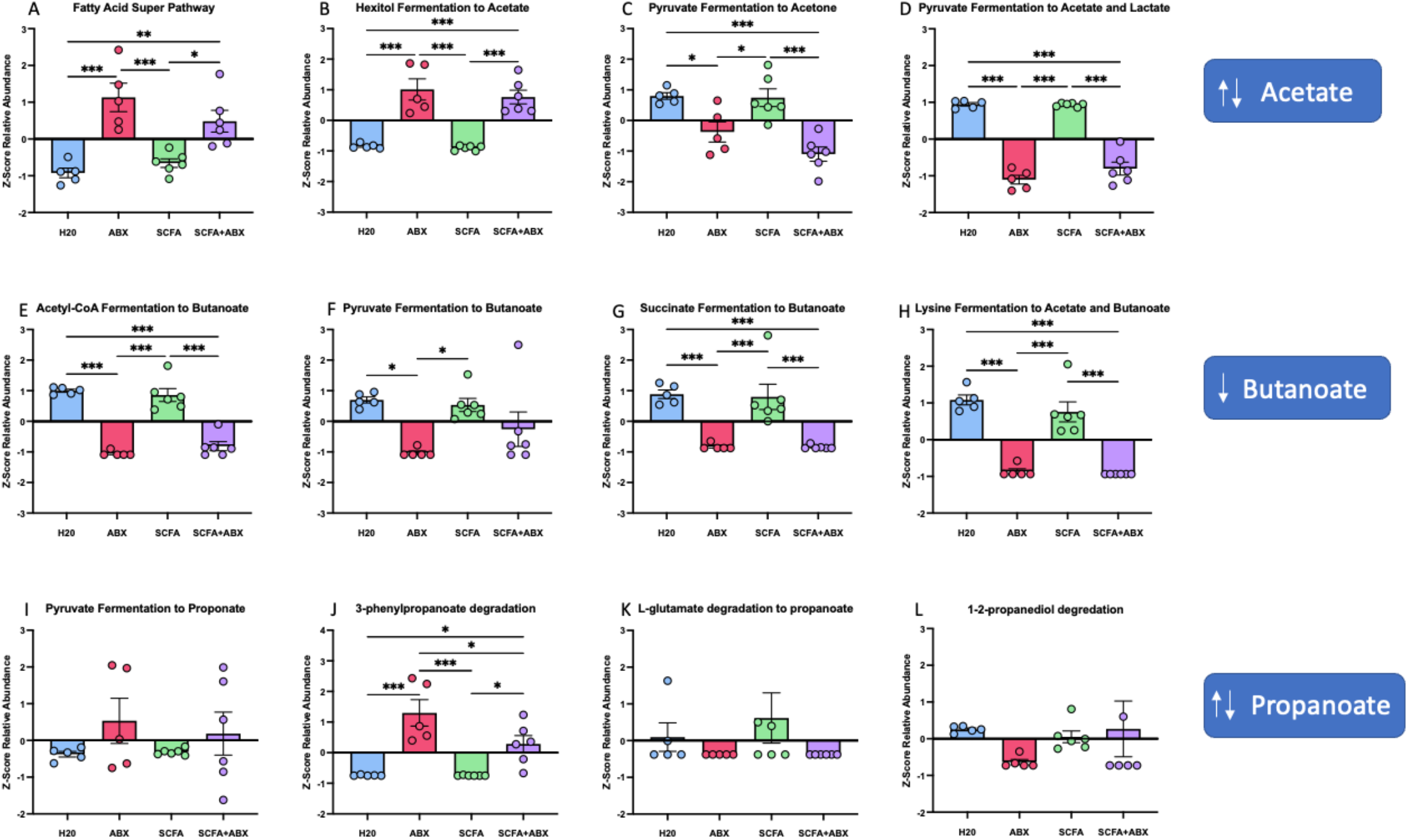
Fatty Acid Pathways are differentially regulated across treatment groups. KEGG Orthologs (KO) generated from PICRUSt2 demonstrate dysregulation of fatty acid metabolism in antibiotic treated animals. Three major fatty acids that are generated by microbiome fermentation of non-digestible fibers are acetate, butyrate (butanoate), and propionate (propanoate). Row 1 demonstrates an increase in FASYN – the super pathway of fatty acid biosynthesis in animals treated with Abx and SCFA+Abx; however, fatty acids are disproportionally increased in the Abx group to produce acetate and propionate (row 3). The Abx group exhibits a significant reduction of butyrate (butanoate) pathways (row 2) compared to the H_2_O, SCFA, and even SCFA+Abx group. This differential regulation of SCFA production and degradation following Abx depletion demonstrates metabolic shifts in microbial communities. \**p*<0.05, \*\**p*<0.01, \*\*\**p*<0.001.

**Supplemental Figure 3.**
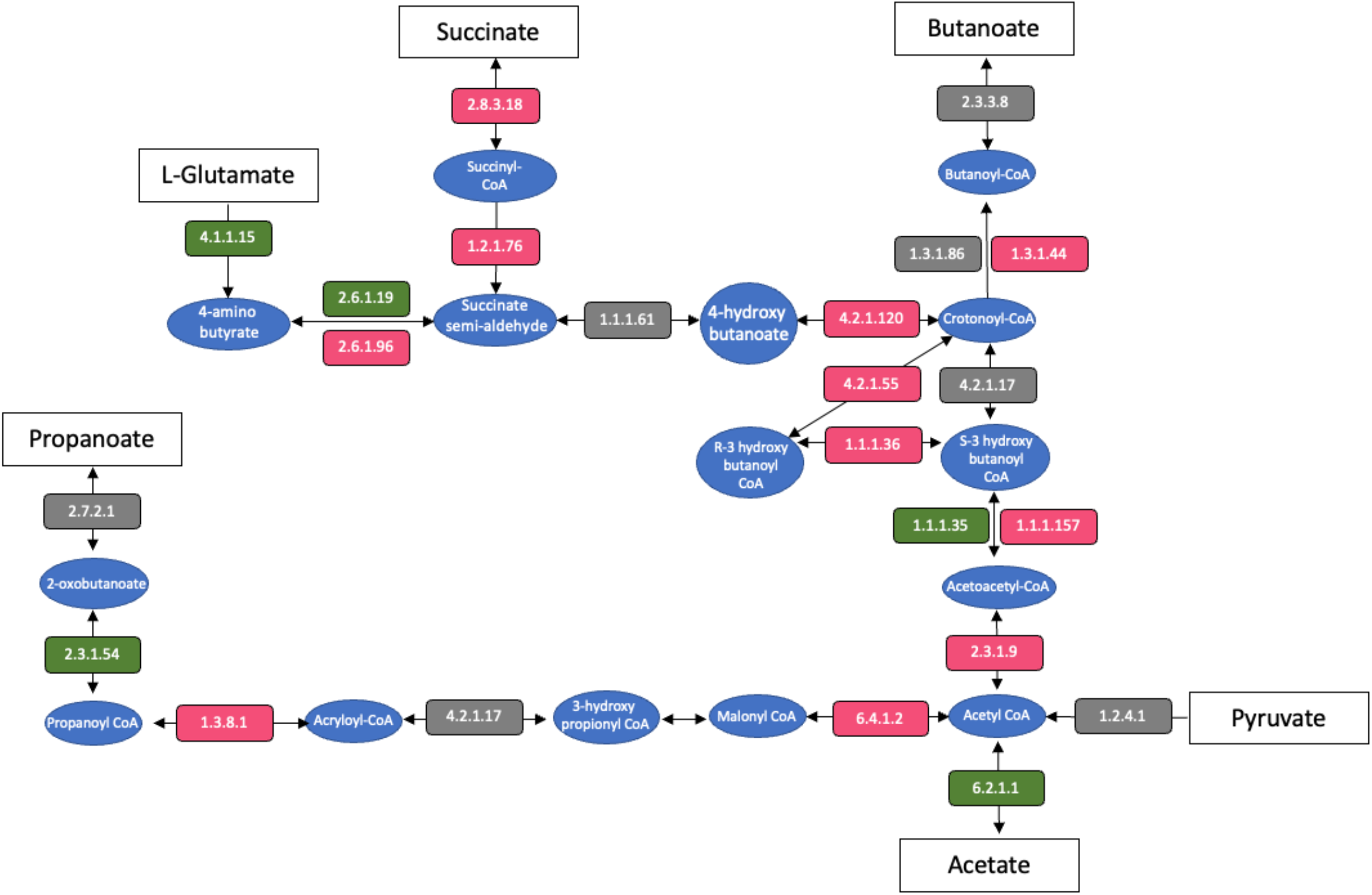
Butanoate production is reduced in the Abx group compared to H_2_O group. MetaCyc enzymatic output from PICRUSt is mapped onto a diagram of fatty acid metabolism. Enzymes that are significantly decreased in the Abx group compared to the H_2_O group are listed in red. Enzymes that are significantly increased following Abx are listed in green, and enzymes that are not significantly increased or decreased are shown in gray. Intermediate compounds are represented as blue ovals.

**Supplemental Figure 4.**
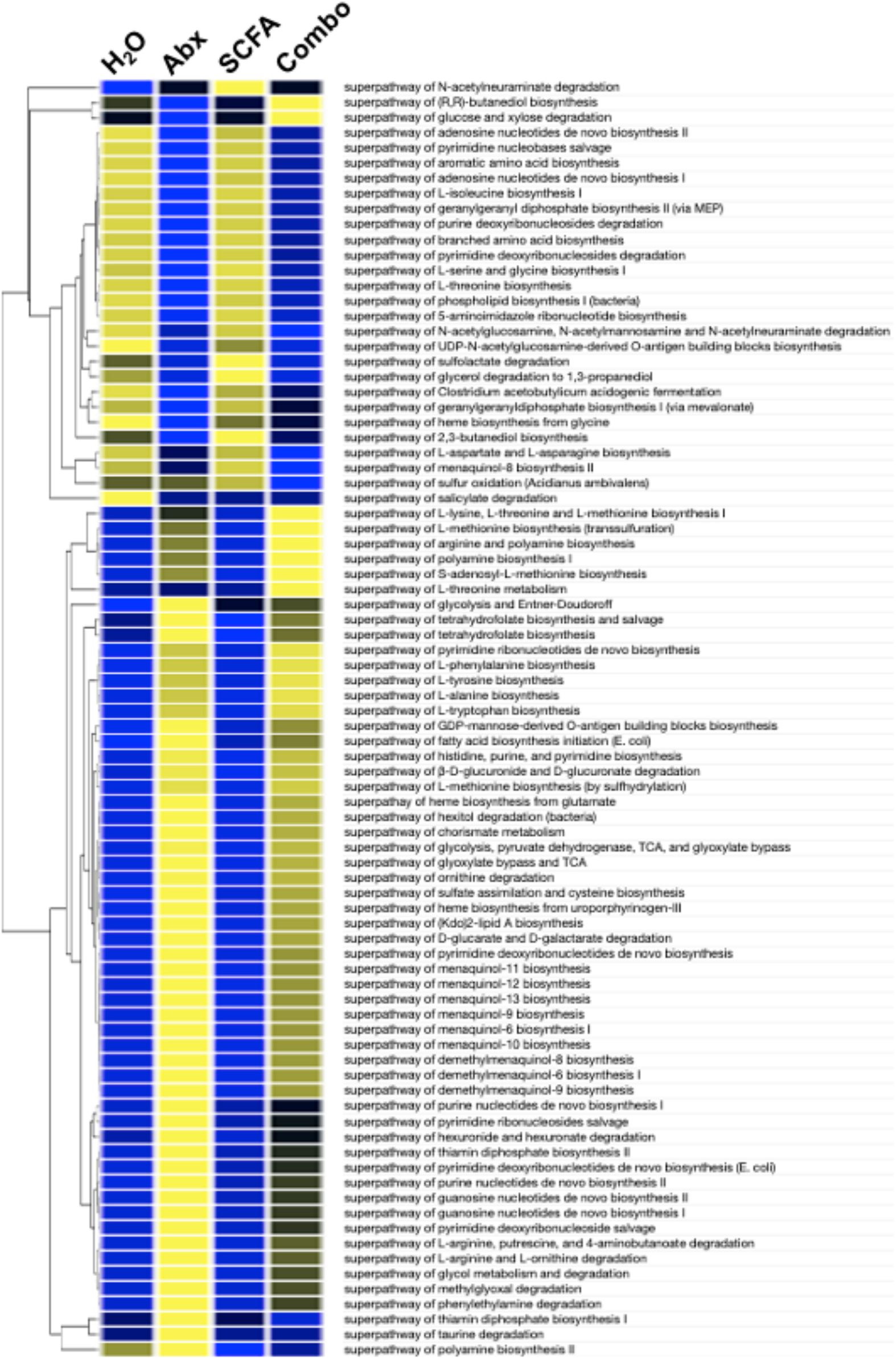
Heat map of KEGG super pathways. Amino acid biosynthesis is decreased in the Abx and Abx+SCFA groups, except for super pathways that are involved in phenylalanine, tyrosine, alanine, and tryptophan biosynthesis which are upregulated in the Abx and Abx+SCFA groups. Menaquinol biosynthesis pathways are highly upregulated in the Abx group and Abx+SCFA groups compared to the H_2_O and SCFA groups.

**Supplemental Figure 5.**
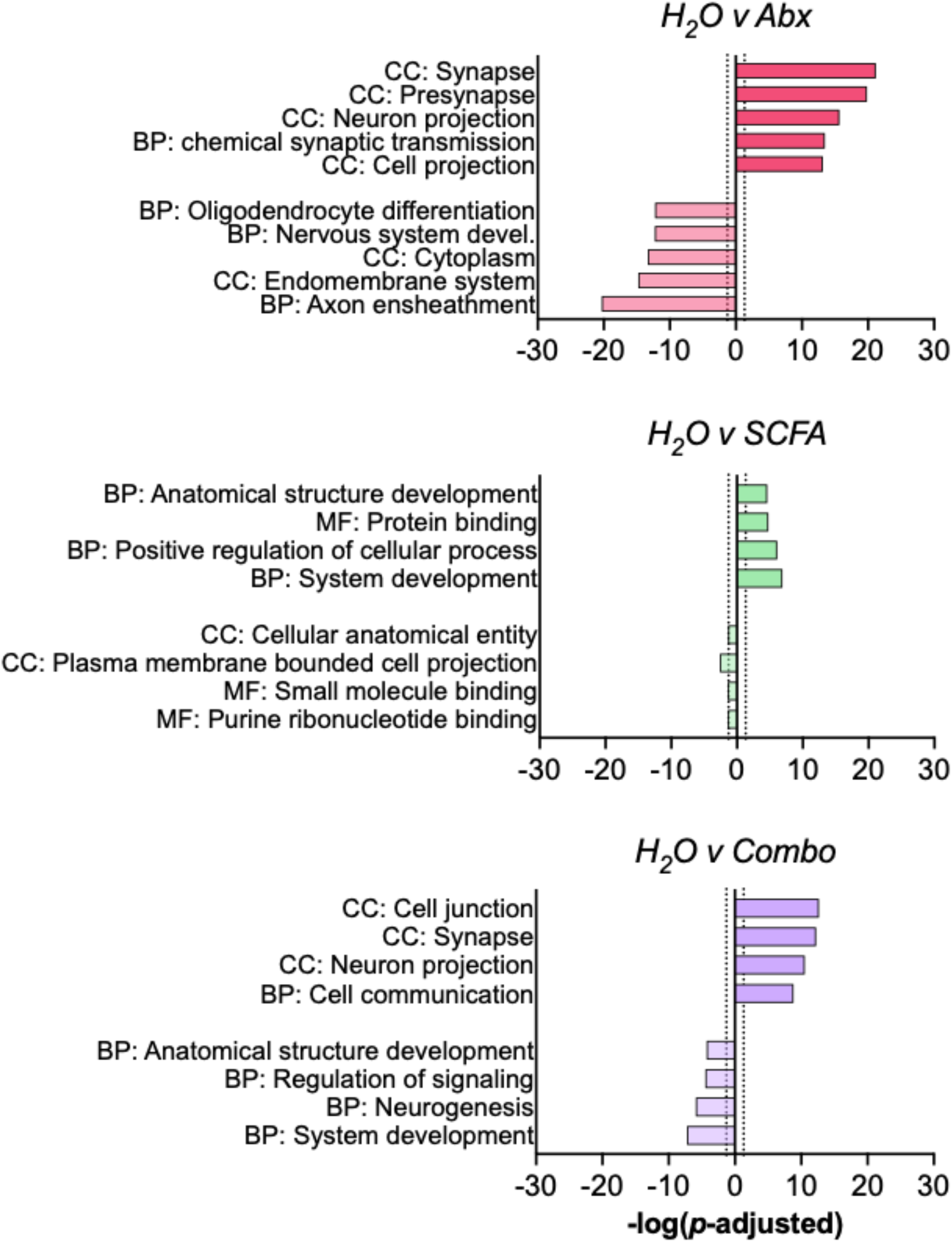
Gene ontologies from RNA-sequencing. Bar graphs showing significant GO terms of interest for each treatment group relative to H_2_O controls. Positive terms are from upregulated genes, and negative terms are from downregulated genes.

